# COMPARISON OF WOODY SPECIES DIVERSITY AND POPULATION STRUCTURE STATUS ACROSS DIFFERENT MANAGEMENT PRACTICE IN THE GOLA NATURAL VEGETATION, EASTER HARARGHE, OROMIA OF ETHIOPIA

**DOI:** 10.1101/2021.09.17.460790

**Authors:** Abdulbasit Hussein, Tasisa Temesgen

## Abstract

The research was carried out at Gola natural vegetation eastern Ethiopia, to identify and Woody species' diversity, richness, evenness, and population structural status will be documented, as well as their diversity, richness, evenness, and population structural status will be analyzed. Using systematic sampling procedures, the woody species diversity and population structure of species were examined in 73 quadrats, each measuring 20 m × 20 m for trees and 5 m × 5 m for shrubs and climbers, within three land use systems: farm land (FL), grazing land (GL), and protected area (PA). The diameter at breast height (DBH), richness, evenness, and density of woody species were all measured in the vegetation. The Shannon Weiner Diversity Index was used to examine the diversity of vegetation. A total of 52 woody species belonging to 33 genera, and there were 24 families found. in Gola natural vegetation. Fabaceae was represented by the highest number of species comprising 8 (18.18 %), 9 (25.00 %) and 8 (32%) of the total number of plant species found in PA, GL, and FL. The PA site had significantly highest population density of vegetation, followed by the GL site, while the FL site had the lowest. The total basal area of PA, GL and FL were 43.73, 31.68 and 22.62 m^2^/ha, respectively. PA site had significantly (P= 0.042) highest Shannon’s diversity index value with mean (3.53) than the others two land use system. This result suggests significance of anthropological disturbance like grazing and farming on woody species diversity and natural forest ecosystem appeared to be adverse dependent on the category and severity of the anthropogenic disturbances.

## Introduction

The richness and evenness (relative abundance) of species among and within living creatures and ecological complexes is referred to as biological diversity (Polyakov et al., 2008). One of the most important analytical aspects of the plant community is the species (Haeussler et al., 2002). Species richness is a straightforward and straightforward metric of biological diversity (Hillebrand et al, 2018). Degradation of vegetation cover and biological diversity due to anthropogenic activities is a rising anxiety in many parts of the world (Hegde and Enters, 2000). The forest coverage in Africa’s is projected to be 650 million hectares, which are 17 percent of the world’s forests counting a several of world biodiversity hotspots (Carney *et al*., 2014). In terms of biological resources (flora and fauna), Ethiopia is recognized as one of Africa's most important countries (Asfaw and Tadesse, 2001). Degradation of forest cover is one of the most pressing issues confronting humanity nowadays (Reynolds et al., 2007). Rapid human population development, agricultural expansion, settlement, poverty, forest cutting, overgrazing, invasive species, and a lack of a strong legislative framework are all key contributors to Ethiopia's forest resource degradation (Lemenih et al., 2014). The primary barrier to total forest conservation in Ethiopia is the lack of inclusion of the local community living surrounding conservation zones into the conservation activities (Woldemariam and Teketay, 2001). The country established a variety of conservation programs, including watershed management, afforestation and reforestation, restoration, and rehabilitation projects, to lessen the damage to natural forests. These methods were developed with the goal of improving vegetation protection and improving the lifestyles of local residents (Mengistu et al., 2005; Wondie, 2015). For several years, the eastern half of Ethiopia has been implementing well-known conservation measures on degraded landscapes, with area exclosure being the most common conservation strategy in this region.

Gola was recognized in 2010 as a protected area, in the east Hararghe zone, East Ethiopia. Gola contains a lot of natural vegetation and a lot of wildlife. However, because it is a freshly constituted protected area, it lacks essential vegetation ecological information. As a result, progressing a comprehensive management plan is critical for effective management and preservation of the area, and this required full baseline information on stand structure and diversity status of woody species, which is essential for long-term management and conservation of the vegetation species. The primary occupations in the Gola natural vegetation region are three types of land use systems: protected natural vegetation by exclosures for more than two decades, agricultural land, and grazing land. Threfore, the study's overall goal was to identify woody species and its diversity with population structure to examine the role of conservation strategy applied in the study area in eastern Ethiopia. A complete floristic analysis of plant species diversity and woody plant population structure was conducted in three land use types: protected natural vegetation, farm land, and grazing land GL, in order to assess the conservation influence status through a complete floristic analysis of plant species diversity and woody plant population structure to fill the current information gap in woody species of Gola natural vegetation.

## 2. Materials and methods

### 2.1. Study area description

Goro Gutu District, also known as the district in the East Hararghe zone, is located in the eastern part of the country. Goro-Gutu, one of the districts in East Hararghe, is part in East Ethiopia. The district covers 531 km2, accounting for approximately 2.35 percent of the zone's total area. Karamile, the capital, is 108 kilometers west of Harar. The agro-climatic zones of Goro-Gutu district are dega (2,000-2,657 m asl), woinadega (1,500-2,000 ms asl), and kola (1,500 m asl), which cover roughly 11, 52 and 37 percent of the district’s total area, respectively. The climatic conditions are defined by the district’s agro-climatic zones. Agriculture (including crop and livestock production) is the people’s primary source of income and employment. In the woreda, the average land holding per household is 0.37 hectares. Mixed farming is used, with sorghum, maize, and wheat being the most prevalent crops grown. The total number of livestock is 195,578. (Zone DPP and FS office, 2001).

The natural vegetation of Gola makes up the majority of the district’s vegetation cover, which was created to protect natural features. The protected area of Gola’s natural vegetation is home to a variety of wildlife and serves as a valuable resource for the indigenous community’s future growth. There is shared grazing pasture (CGL) and farm land used by the local people adjacent to the protected area of Gola natural vegetation, and access to these resources is unrestricted.

### 2.2. Sampling Design

The following three categories of land cover were investigated: (1) protected area (known as “non-arable and settlement”); (2) farm land (with different crops); (3) communal grazing land (‘pasture). Systematic sampling design was carried out to gather data on the vegetation of woody species at the study location, they are dissimilar in their vegetation distribution pattern and types. The whole area of the study site was used to create appropriate transect lines and sampling quadrats for vegetation data collection in each of the land use groups. Total of six transect lines were laid (two in each land cover type). To avoid the effect of disturbances the first and the last transects lines were laid at a distance of 50 m from the edges. Along the transect lines, a total of 68 systematically selected study quadrats (20 from PA, 28 from Arab land and 25 from CGL). At every 100 m interval, quadrats of 20m×20m (400m2) were systematically created. To collect data on shrubs and climbers, five 5m×5m (25m2) sub quadrats were employed and averaged at the four corners and center of the main quadrats.

### 2.3. Data collection methods

In each land cover type entire the woody species vegetation encountered in all quadrats were noted and coded with scientific and local namesWith a hypsometer and a clipper, woody vegetation with DBH >2.5 cm was measured and recorded for height and diameter at breast height (DBH) respectively in each quadrat. Individuals with a single stem that branched over breast height were classified as trees. Shrubs with many straight trunks perceptibly related at ground level or one solo trunk with attached branches under breast height were smaller than ten meters in height (Powell, 2005; Nzunda, et al. 2007). The circumference of branching trees and bushes around breast height was measured separately and averaged.

#### 2.3.1 Breast Height Diameter (DBH)

A clipper was used to measure DBH at a distance of roughly 1.3 m from the ground. Circumference at breast height was measured and recorded for woody species such as trees and shrubs with DBH > 2.5 cm (DBH). Individual trees/shrubs with several stems or forks below 1.3 m in height were also evaluated (Kent and Coker, 1992). The diameter of trees and bushes branching at the breast height was measured separately and averaged.

#### 2.3.2. Height

Height is a simple parameter that can be used for direct measurement. The heights of the tree were measured using Hypsometer. Identification vegetation species was done on the study sites. Specimens were prepared and taken to Haramaya University’s herbarium for identification of species that were not identified in the field.

### 2.4. Analysis

Diversity has been the most often utilized criterion for evaluating a site’s conservation potential and ecological significance (Magurran, 1988). The best and most extensively used diversity index is the Shannon diversity index. It takes into account both the diversity and evenness of species in a group. This measure takes species composition and evenness within a given territory or community into account. To examine the level of species diversity and evenness of species distribution, the Shannon diversity indices of diversity and evenness were applied (Kent and Coker, 1992)

The following is how the Shannon diversity index was determined.

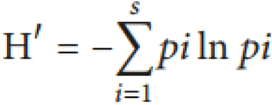

Species richness was calculated using all of the species found in each plot.

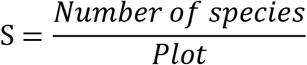

Evenness or equitability is a measure of how closely the abundances of different species in a given location are identical (Krebs, 1999; Magurran, 2004). The relative abundance of the different species that make up an area’s richness is measured by species evenness (measure of species balance).

The following formula is used to calculate evenness: 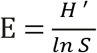 Where E stands for evenness, H’ for Shannon-Wiener diversity index, and S stands for species.

The crowdedness of woody species diameter at breast height or stamp height (DBH/DSH) is known as the basal area (BA). It’s a measure of dominance and a way to see the cross-section of a forest stand, and it’s calculated using the formula below (Kent and Coker, 1992):

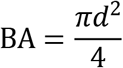

Where BA = basal area in square meters per hectare, d = breast/stump height diameter in meters, and π= 3.14

The quantity of plants of a particular species per unit area is known as density. It’s linked to abundance, but it’s more effective for determining a species’ importance. The sum of individuals per species was determined in terms of species density per convenient area unit such as a hectare, using a small quadrat inserted many times into the vegetation communities under study (Mueller-Dombois and Ellenberg, 1974).

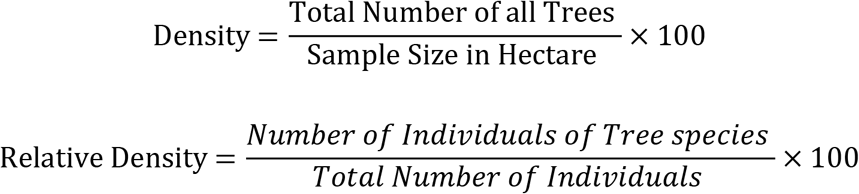

The probability or chance of finding a plant species in a given sample area or quadrat is defined as the number of quadrats occupied by a given species per number thrown or, more commonly, as a percentage. Frequency was calculated using quadrats and expressed as the number of quadrats occupied by a given species per number thrown or, more often, as a percentage. It is calculated as follow.

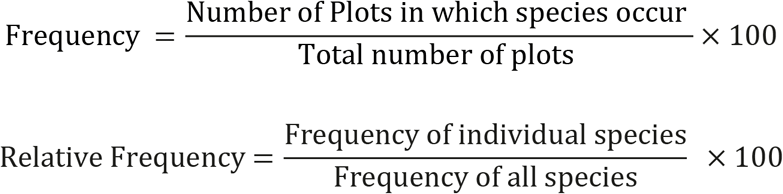

Relative dominance is the ratio of dominance of individual tree species per dominance of all tree species.

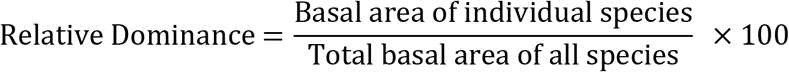

The ecological value of woody species was compared using Importance Value Indices (IVI) based on three criteria (relative frequency, relative density, and relative abundance). The significance of value index (IVI) was generated for each woody species using the formula below.

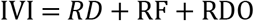

The Jackard similarity index was used to determine the similarity coefficient between land cover types. The pattern of species turnover among the different community types was determined using Jaccard’s similarity index. This is how it was calculated: (Chidumayo, 1997).

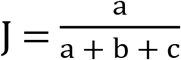

Where J is the Jaccard similarity coefficient, a is the number of species that are found in both samples, b is the number of species that are only found in the first site, and c is the number of species that are only found in the second site. To get a percentage similarity index, the coefficient is often multiplied by 100.

All individuals encountered in the quadrats were divided into height and diameter classes to examine the population structure of woody plants (Emiru et al., 2002).

Following data collection, data was coded and arranged for analysis. Descriptive and inferential statistics were employed to analyze the quantitative data. The proportion of structural parameters of vegetation was discussed using descriptive statistics such as mean, frequency, density, and percentage. In addition to this, inferential statistics such as one-way analysis of variance (ANOVA) was employed. ANOVA was attributed to data that generated from vegetation (species richness, evenness and Shannon-Weiner Diversity Index) to examine the differences among the three-land use type. Statistical analyses were conducted using R software program (version 3.5.1.) using vegan, lattice, permute and biodiversityR package (R Core Team, 2018) at 5% significance level. Tables and Figures were used to present the result of descriptive and inferential statistics of the study.

## 3. Results and discussion

### 3.1. Composition of woody species

A total of 52 woody species belonging to 33 genera and 24 families were collected in Gola natural vegetation. These species growth form distribution was 23 (44.23 %) shrubs, 16 (30.77 %) trees, 7 (13.46 %) climbers, and 6(11.54%) tree/shrubs. Among these, 44 species of plants were collected from protected area site, of which 19 (43.18%) were shrub species, tree accounted 14(31.82%) whereas climbers and tree/shrubs take a minimal amount 7(15.91%) and 4(9.09%) respectively. Like PA in GL the proportion of shrub took the paramount proportion with 17(47.25 %), followed by tree 12 (33.33%) and tree/shrubs and climbers share 4(11.11%) and 3(8.33%) respectively. Similarly, in HD site the proportion of shrubs possessing 13(52) %, followed by trees 9(36%) and, tree/shrubs and climber share the lowest value 2(8%) and 1(4%) respectively.

Overall, PA had a larger abundance and dispersion of plant species than GL and FL. The difference in woody species composition between the three land cover categories revealed that adequate conservation techniques such as area enclosures and restoration practices had a progressive influence on vegetation maintenance. The decrease in woody species variety in grazing land, according to Kibret (2008), could be a symptom of the vegetation species’ increased susceptibility to animals and/or anthropogenic interference at maturity or early stages of regeneration. This could indicate that nearby residents and/or their domestic animals collected persons in the GL and FL at an early growth stage (Wondie et al., 2014). Similarly, Sisay et al. (2001) and Tessema et al. (2011) hypothesized that extensive grazing/browsing could result in a decrease in plant species density and diversity over time.

Fabaceae was found to be the most species-rich family, accounting for 8 (18.18%), 9 (25.00%), and 8 (32%) of the total plant species identified in PA, GL, and FL, respectively. Euphorbiaceae was found to be the second most species-rich family, accounting for 4 (16.00%) and 4 (11.11%) of the total plant species identified in FL and GL, respectively. However, Euphorbiaceae and Oleaceae were the second most species-rich family in Pennsylvania, with 5 (11.36 percent) of species. Similar investigations in the Gra-Kahsu forest (Atsbha et al., 2019) and Hugumburda forest (Aynekulu, 2011) found that the Fabaceae family dominated the vegetation stands. Similarly, the Fabaceae family dominated the Babile Elephant Sanctuary (Anteneh et al., 2011). This dominance of the Fabaceae family could be attributed to the family’s ability to adapt to the country’s different ecological conditions.

### 3.2. Vegetation Structure

#### 3.2.1. Species Density of Woody Plants

The number of plants per quadrat area is expressed as density, and it is a critical measure for long-term forest management. The average overall density of Gola natural vegetation was estimated around 8844.16 individuals per hectare. In Gola Natural Vegetation, the six most abundant woody species are listed in order of density were *Acacia Senegal, Grewia schweinfurthii, Lantana camara, Acalypha fruticosa and Grewia erythraea.*

The highest density of single woody species of *Acacia Senegal* accounted a total of 2240 and 960 number of plants per hectare in PA and GL respectively whereas, *Grewia schweinfurthii* 420 individuals per hectare in the FL. The woody vegetation **s**pecies in the grazing land continued maybe due to they had a tolerance to anthropogenic disturbance and hereafter, are very significant in the restoration of degraded natural vegetation in the study area. Likewise, the maximum densities of some species like *L. camara,* could be because of their unpalatable characteristics for both domestic and wild animals and extensive range of seed dispersal system and highly reproductive strategies (Feyera, 2006; Anteneh *et al.,* 2011). Alike study also stated that open-grazed lands had a fewer vegetation density as associated to area exclosures (Teshome *et al*., 2009; Yayneshet, 2011).

#### 3.2.2. Frequency

Frequency is defined as the number of quadrats (expressed as a percentage) in which a specific species was discovered in the research area. The uniformity of forest composition can be seen in the smaller number of species in lower frequency classes and the higher number of species in higher frequency classes. And the higher number of species in love frequency classes and the low number of species in higher frequency classes shows heterogeneity of species. In these study woody vegetation species were clustered into four frequency classes: A ≤25; B = 26– 50; C = 51–75; D ≥75. The current study found a high percentage of woody species in lower frequency groups and a low percentage of species in higher frequency classes. This indicates that generally the study sites had heterogeneous species composition. PA had more species percent with higher frequency class (class D) which are 11.33% as compare to GL and FL that had only 5.5% and 4% of species respectively. While in lower frequency class (A) GL had species percent (58.33%) than FL (52%) and PA (50%) (Fig 2). As a result, the study confirms that each land use system has a significant degree of floristic heterogeneity.

**Figure 1.**
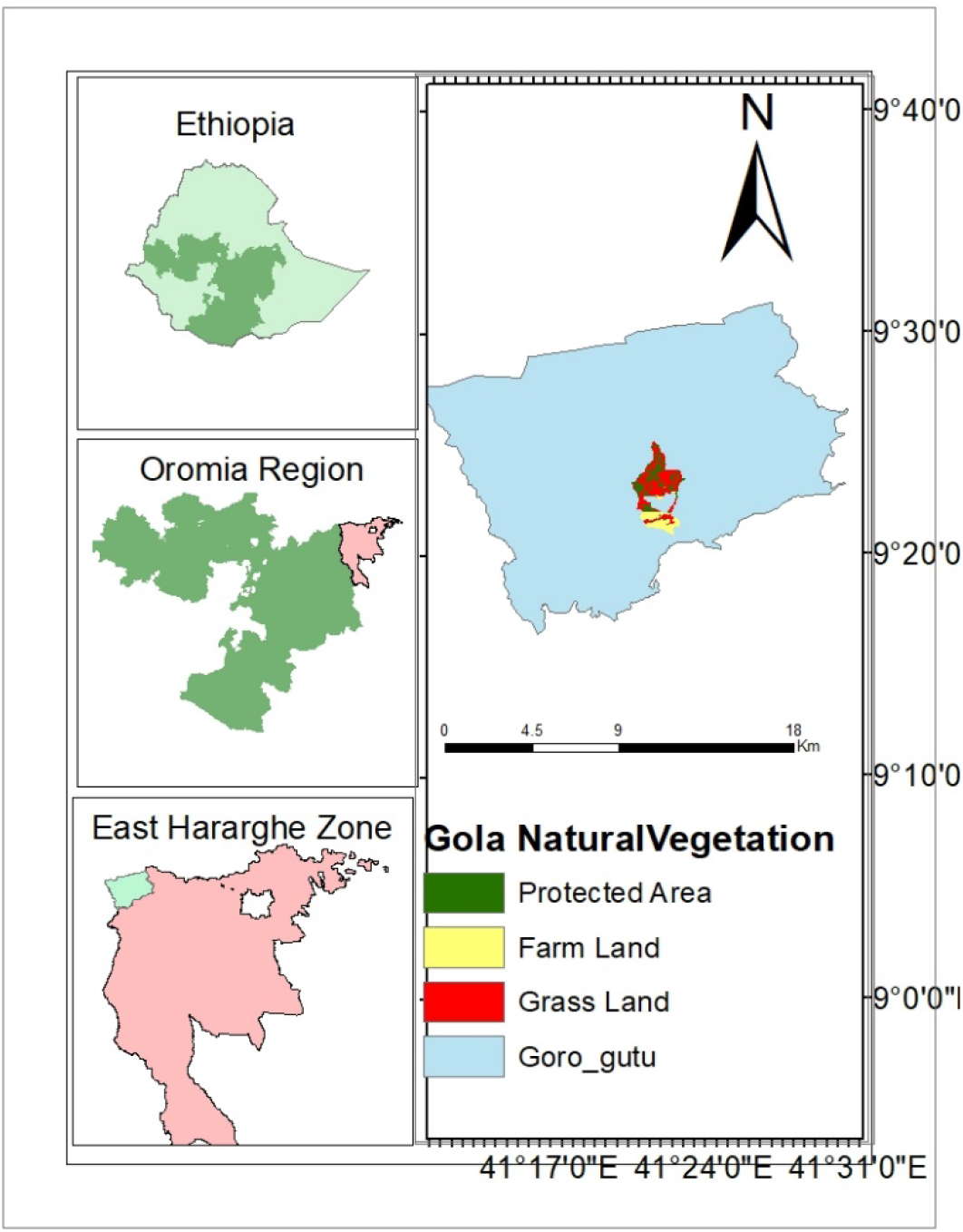
Location map of Gola natural vegetation elephant sanctuary

**Figure 2.**
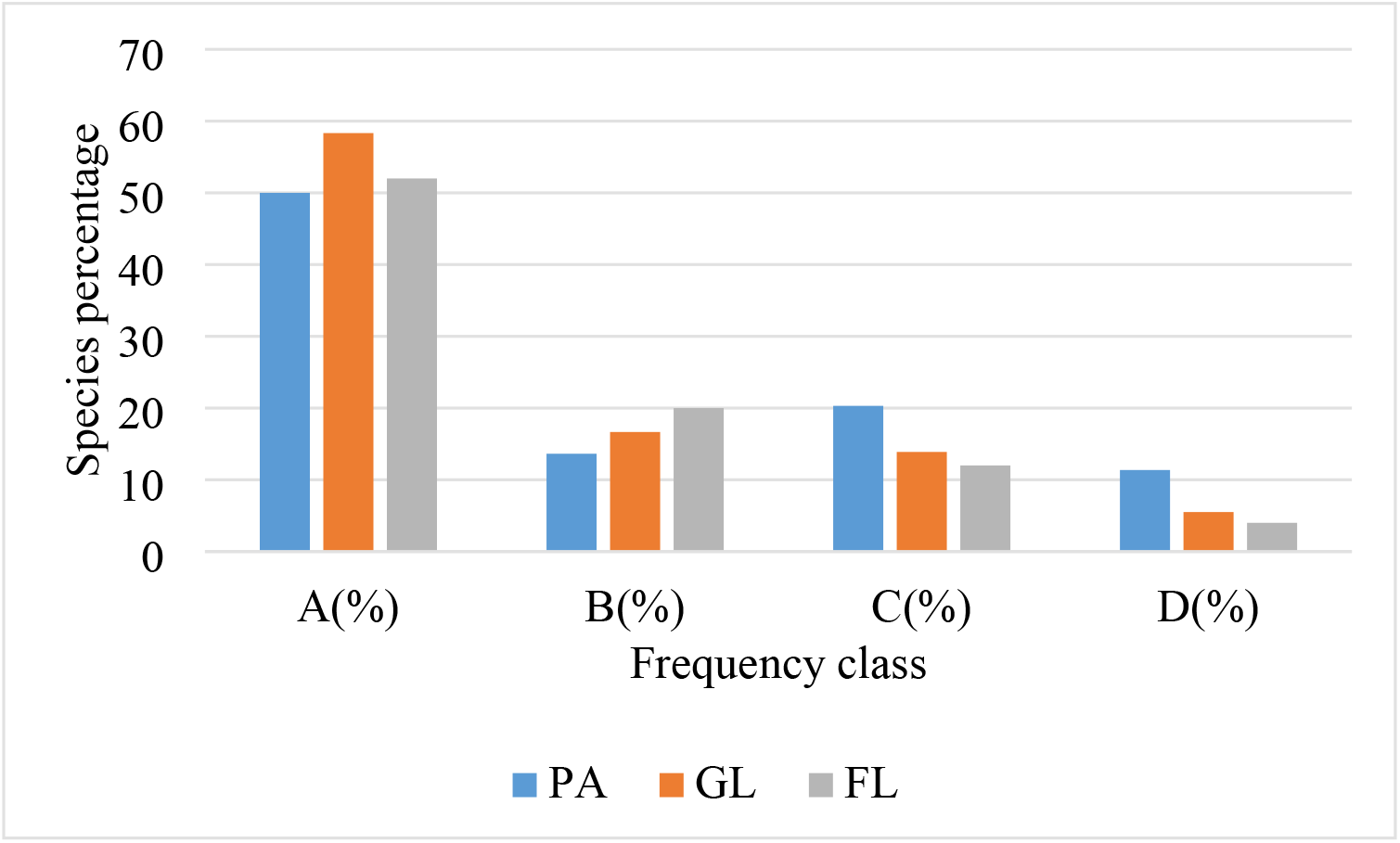
Woody species frequency class distribution at all three land use sites Classifications of frequencies: A ≤25; B = 26– 50; C = 51 – 75; D ≥75 (PA= protected area, GL=grazing land and FL=farm land)

*Acacia bussei* was the most frequently recorded woody plant species in both PA and GL, whereas *Grewia schweinfurthii* have been the most frequently recorded in FL. However, *Olea europaea, Grewia velutina and Sansevieria ehrenbergii* were the least frequently recorded woody plant species in PA, GL and FL respectively, each were recorded only in one plot. The reason for lack of perfectly frequent for some species in the Gola natural vegetation could be due to relatively higher human intervention and animal grazing. Species of such least frequency need to be given priority in conservation to enhance their frequency distribution.

### 3.3. Diversity, Richness, and Evenness of Woody Species

The mean diversity and evenness of woody species were 3.17 and 0.85 respectively in Gola natural vegetation. Gola natural vegetation had medium Shannon diversity index. However, the three land use systems had significantly different value of species diversity and values of evenness. The Shannon diversity index value of PA was significantly greater than that of FL. However, the GL had not significantly different from each other, even though GL slightly higher than FL (Table 1). This could be the result of frequent habitat disruptions in the GL and FL as a result of daily and intensive human and livestock activity for grazing and other agricultural purposes. The inability of seedlings of woody species to establish at an early stage of growth, as well as selective defoliation and crushing by grazing animals, may be contributing to the loss of species diversity in the GL and FL (Belaynesh, 2006).

**Table 1.** Mean ± SD of species diversity, richness and evenness of Gola natural vegetation.

| Land use types | Shannon diversity index ($H'$) | Evenness ($H'/H_{max}$) | Species richness |
| --- | --- | --- | --- |
| PA | 3.53 <sup>a</sup>$\pm 0.82$ | 0.91 <sup>a</sup>$\pm 0.023$ | 19.7<sup>a</sup>$\pm 7.50$ |
| GL | 2.98 <sup>ab</sup>$\pm 0.75$ | 0.86<sup>b</sup>$\pm 0.03$ | 14.3 <sup>a</sup>$\pm 7.63$ |
| FL | 2.21 <sup>b</sup>$\pm 0.63$ | 0.63<sup>b</sup>$\pm 0.06$ | 11.2 <sup>a</sup>$\pm 7.83$ |
| P-value | 0.042* | 0.047 * | 0.80 |

Different letters indicate the significance differences.
\* indicates significant at 5 % significance level
(PA= protected area, GL=grazing land and FL=farm land)

In three land use types mean evenness decreased significantly (p=0.04) from the PA site through the GL to the FL site. A low mean evenness rating in FL suggests that only a few species dominate in the community. While high evenness means in PA indicates, there are constant dispersals of the species in a given ecological community (Cavalcanti and Larrazabal, 2004).

Mean of richness of woody species decreased non significantly from the PA site through the GL to the FL land use types shows average number of species per sampling unit was also higher in the PA than in the GL and FL land use type in Gola natural vegetation (Table 1)

**Table 2.** Similarity of woody species between three land use system

| Land use type | PA | GL | FL |
| --- | --- | --- | --- |
| PA | 0 |  |  |
| GL | 0.78 | 0 |  |
| FL | 0.52 | 0.65 | 0 |

(PA= protected area, GL=grazing land and FL=farm land)

### 3.4 Jaccard Coefficient of Species Similarity

The similarity in family, genus, and species composition of different sites of the three land use groups was calculated using Jaccard’s coefficient of similarity. The Jaccard’s similarities of woody species for PA and GL were 78% of species. Similarly, 65%, similarity values were obtained from GL and FL land use types. However, the similarities between HD and LD were found to be 52% of species. Highest similarity value was obtained for the species level between PA and GL followed by GL and FL sites, while the lowest were recorded in between PA and FL.

### 3.5. Basal area

The cross-sectional area of all the stems in a stand at breast height is known as the basal area (BA) (1.3 m above ground level). For woody species with DBH > 2.5 cm, the mean basal area (reported as the basal area of stems per hectare) in Gola Natural vegetation was 33.11 m2 ha-1. The total basal area for PA, GL and FD were 43.73 m^2^/ha, 31.68 m^2^/ha and 22.68 m^2^/ha, respectively. PA had significantly higher basal area coverage than the other two sites and followed by GL (P < 0.05). This maybe be due to the existence of relatively higher percentage of larger and matured trees leading to bigger diameters in PA because of slight human interference activities like logging of trees, farming practice and animal grazing. But FL had significantly lower basal area coverage. This directly relational trend indicated that human intervention influences basal area of vegetation.

The most important species in the vegetative community could be those having the highest basal area. The highest proportion of mean BA at the PA was covered by *Acacia seyal* (4.52 m^2^/ha), followed by *Euphorbia adjurana* (3.79 m^2^/ha). While, *Acacia bussei* (3.62 m^2^/ha) and *Acacia senegal* (3.14 m^2^/ha) were accounted the highest proportion of the mean BA and followed by *Acacia tortilis* (3.46 m^2^/ha) and *Acacia bussei* (2.54 m^2^/ha) at GL and FL respectively. In this study, cross-sectional assessment of individual species in the basal area revealed that only a few or small woody species dominated the studied area. This also shows that species with the biggest basal area do not always have the highest density, proving that species differ in size (Ayanaw and Gemado, 2018).

### 3.6. Importance value index

The IVI values have aided in recognizing a species’ structural relevance in community organization (Premavani et al., 2014). It’s also used to compare the ecological significance of species, with a high IVI score indicating a high social structure in the community. According to Shibru and Balcha (2004), species with the greatest IVI are the dominant species in a given vegetation. The IVI of the vegetation’s woody species was determined using three structural characteristics (Relative Dominance, Relative Frequency, and Relative Density). In this respect, the results of IVI in PA revealed that for *Acacia Senegal* (27.98), *Grewia schweinfurthii* (25.77) and *Lantana camara* (16.37) were the three species with higher important value index. These woody plants could be ecologically essential in the natural vegetation of Gola’s PA. Whereas, woody species like, *Jasminum abyssinicum* (0.67*), Jasminum grandiflorum* (0.68) and *Jasminum abyssinicum* (0.72) found with lower IVI (Table 3). This indicates that these species are the least ecologically significant species in the site. Their lower IVI may signal that these woody species are imperiled and require prompt conservation measures, among other things (Anteneh et al., 2011; Temesgen et al., 2015).

**Table 3.** Woody species of the Gola natural vegetation’s relative frequency, density, dominancy, and important value index in three land use systems.

| Species name | PA |  |  |  | GL |  |  |  | FL |  |  |  |
| --- | --- | --- | --- | --- | --- | --- | --- | --- | --- | --- | --- | --- |
|  | RF | RD | RDO | IVI | RF | RD | RDO | IVI | RF | RD | RDO | IVI |
| *Acacia brevispica* | 2.85 | 2.27 | 0.25 | 5.39 | 5.44 | 10.9 | 0.19 | 16.5 | 2.54 | 4.61 | 2.74 | 9.8 |
| *Acacia bussei* | 6.66 | 1.23 | 3.07 | 10.9 | 6.93 | 1.14 | 11.4 | 19.5 | 7.52 | 0.98 | 11.2 | 19.7 |
| *Acacia etbaica* | 5.23 | 0.46 | 5.18 | 10.8 | 4.45 | 0.55 | 8.02 | 13.0 | 7.52 | 0.53 | 7.78 | 15.8 |
| *Acacia negrii* | 4.28 | 5.08 | 0.31 | 9.68 | 5.44 | 9.39 | 0.37 | 15.2 | 5.83 | 11.2 | 4.98 | 22.0 |
| *Acacia nilotica* | 4.76 | 0.36 | 4.59 | 9.71 | 3.96 | 0.34 | 8.94 | 13.2 | 6.66 | 0.49 | 4.98 | 12.1 |
| *Acacia senegal* | 6.19 | 19.62 | 2.17 | 27.9 | 6.43 | 12.1 | 0.43 | 19.0 | 4.16 | 5.27 | 13.8 | 23.2 |
| *Acacia seyal* | 4.28 | 0.28 | 10.3 | 14.9 | 4.95 | 0.44 | 9.91 | 15.3 | 0 | 0 | 0 | 0 |
| *Acacia tortilis* | 3.33 | 1.56 | 5.81 | 10.7 | 5.94 | 0.93 | 10.9 | 17.8 | 5.83 | 0.78 | 6.78 | 13.3 |
| *Acalypha fruticosa* | 4.76 | 5.95 | 0.06 | 10.7 | 4.95 | 7.36 | 0.20 | 12.5 | 7.50 | 11.2 | 0.37 | 19.0 |
| *Agave sisalana* | 0 | 0 | 0 | 0 | 0 | 0 | 0 | 0 | 2.50 | 3.29 | 0.18 | 5.98 |
| *Aloe harlana* | 1.42 | 1.40 | 0.50 | 3.33 | 2.97 | 3.55 | 0.50 | 7.02 | 0 | 0 | 0 | 0 |
| *Balanites glabra* | 3.80 | 3.68 | 0.58 | 8.07 | 4.45 | 9.14 | 0.61 | 14.2 | 0 | 0 | 0 | 0 |
| *Barbeya oleoides* | 0 | 0 | 0 | 0 | 0.49 | 0.25 | 0.00 | 0.75 | 0 | 0 | 0 | 0 |
| *Berchemia discolor* | 1.90 | 0.09 | 5.18 | 7.19 | 0.99 | 0.06 | 8.02 | 9.08 | 0 | 0 | 0 | 0 |
| *Bridelia micrantha* | 0.95 | 0.03 | 4.59 | 5.57 | 0.99 | 0.03 | 7.58 | 8.60 | 0 | 0 | 0 | 0 |
| *Cadia purpurea* | 2.85 | 3.50 | 0.20 | 6.56 | 1.48 | 2.54 | 0.22 | 4.24 | 3.33 | 4.61 | 0.25 | 8.20 |
| *Clerodendrum* |  |  |  |  |  |  |  |  |  |  |  |  |
| *cephalanthum* | 0 | 0 | 0 | 0 | 0 | 0 | 0 | 0 | 1.66 | 2.63 | 2.50 | 6.80 |
| *Dodonaea angustifolia* | 3.33 | 2.97 | 0.20 | 6.51 | 1.98 | 1.77 | 1.79 | 5.54 | 1.66 | 1.97 | 0.63 | 4.28 |
| *Dracaena ombet* | 0 | 0 | 0 | 0 | 0 | 0 | 0 | 0 | 0.83 | 0.04 | 0.02 | 0.89 |
| *Dregea rubicunda* | 0.95 | 0.70 | 0.05 | 1.71 | 0 | 0 | 0 | 0 | 0 | 0 | 0 | 0 |
| *Euphorbia abyssinica* | 0 | 0 | 0 | 0 | 0 | 0 | 0 | 0 | 1.66 | 0.16 | 3.32 | 5.15 |
| *Euphorbia adjurana* | 1.42 | 1.57 | 8.68 | 11.6 | 1.48 | 1.77 | 2.14 | 5.40 | 2.50 | 1.97 | 2.74 | 7.21 |
| *Euphorbia candelabrum* | 0.95 | 0.03 | 7.54 | 8.52 | 1.48 | 0.07 | 5.57 | 7.13 | 0.83 | 0.04 | 11.2 | 12.0 |
| *Euphorbia polyacantha* | 0.95 | 0.87 | 0.07 | 1.90 | 0 | 0 | 0 | 0 | 0 | 0 | 0 | 0 |
| *Ficus vasta* | 1.90 | 0.09 | 7.17 | 9.18 | 3.96 | 0.30 | 7.16 | 11.4 | 0 | 0 | 0 | 0 |
| *Grewia erythraea* | 2.85 | 3.85 | 0.08 | 6.79 | 1.98 | 3.04 | 0.47 | 5.50 | 5.00 | 6.59 | 0.40 | 11.9 |
| *Grewia ferruginea* | 0.47 | 0.35 | 0.09 | 0.91 | 0 | 0 | 0 | 0 | 1.66 | 1.97 | 4.18 | 7.83 |
| *Grewia schweinfurthii* | 6.19 | 19.10 | 0.48 | 25.7 | 4.45 | 7.36 | 0.25 | 12.0 | 10.8 | 13.8 | 0.31 | 24.9 |
| *Grewia tenax* | 1.90 | 1.57 | 0.64 | 4.12 | 0 | 0 | 0 | 0 | 0 | 0 | 0 | 0 |
| *Grewia velutina* | 0 | 0 | 0 | 0 | 0.49 | 0.25 | 0.26 | 1.01 | 0 | 0 | 0 | 0 |
| *Grewia villosa* | 0.95 | 0.52 | 0.18 | 1.66 | 0 | 0 | 0 | 0 | 0.83 | 0.65 | 0.06 | 1.56 |
| *Jasminum abyssinicum* | 0.47 | 0.17 | 0.07 | 0.72 | 0 | 0 | 0 | 0 | 0 | 0 | 0 | 0 |
| *Jasminum abyssinicum* | 0.47 | 0.17 | 0.02 | 0.67 | 0.49 | 0.25 | 0.05 | 0.80 | 0 | 0 | 0 | 0 |
| *Jasminum grandiflorum* | 0.47 | 0.17 | 0.03 | 0.68 | 0 | 0 | 0 | 0 | 0 | 0 | 0 | 0 |
| *Jasminum schimperi* | 0.47 | 0.35 | 0.04 | 0.86 | 0.49 | 0.50 | 0.06 | 1.07 | 0 | 0 | 0 | 0 |
| *Lantana camara* | 5.23 | 10.8 | 0.27 | 16.3 | 6.93 | 9.39 | 0.20 | 16.5 | 6.66 | 12.5 | 0.16 | 19.3 |
| *Olea europaea* | 0.47 | 0.01 | 5.81 | 6.30 | 0 | 0 | 0 | 0 | 0 | 0 | 0 | 0 |
| *Opuntia ficus-indica* | 3.33 | 3.32 | 0.15 | 6.81 | 1.98 | 2.28 | 0.19 | 4.46 | 5.83 | 7.25 | 0.21 | 13.30 |
| *Opuntia stricta* | 2.85 | 1.92 | 0.27 | 5.05 | 0.99 | 1.27 | 0.19 | 2.45 | 0 | 0 | 0 | 0 |
| *Pittosporum abyssinicum* | 1.42 | 1.05 | 0.36 | 2.84 | 1.48 | 1.01 | 0.37 | 2.87 | 1.66 | 1.97 | 5.40 | 9.05 |
| *Rhoicissus tridentata* | 0.95 | 0.35 | 0.07 | 1.37 | 0.99 | 0.76 | 0.08 | 1.83 | 0 | 0 | 0 | 0 |
| *Rhus glutinosa* | 1.42 | 0.07 | 6.47 | 7.98 | 0.99 | 0.04 | 7.16 | 8.19 | 0 | 0 | 0 | 0 |
| *Rhus natalensis* | 1.90 | 1.22 | 0.58 | 3.71 | 3.46 | 4.31 | 0.33 | 8.12 | 0 | 0 | 0 | 0 |
| *Rhus ruspolii* | 0.47 | 0.02 | 7.91 | 8.41 | 0 | 0 | 0 | 0 | 0 | 0 | 0 | 0 |
| *Rhus vulgaris* | 0.47 | 0.03 | 7.91 | 8.42 | 0.49 | 0.01 | 4.85 | 5.36 | 0.83 | 0.65 | 8.85 | 10.3 |
| *Rumex nervosus* | 0 | 0 | 0 | 0 | 1.48 | 0.76 | 0.25 | 2.50 | 0 | 0 | 0 | 0 |
| *Salvadora persica* | 0.95 | 1.05 | 0.66 | 2.67 | 2.47 | 4.06 | 0.69 | 7.23 | 3.33 | 3.95 | 6.78 | 14.0 |
| *Sansevieria ehrenbergii* | 0.95 | 0.52 | 0.05 | 1.53 | 0 | 0 | 0 | 0 | 0.83 | 0.65 | 0.00 | 1.49 |
| *Solanum nigrum* | 0.95 | 0.70 | 0.41 | 2.06 | 0 | 0 | 0 | 0 | 0 | 0 | 0 | 0 |
| Species | RF | RD | RDO | IVI | RF | RD | RDO | IVI | RF | RD | RDO | IVI |
| *Withania somnifera* | 0 | 0 | 0 | 0 | 0.49 | 0.25 | 0.03 | 0.78 | 0 | 0 | 0 | 0 |
| *Ziziphus mauritiana* | 0.47 | 0.17 | 0.28 | 0.93 | 0 | 0 | 0 | 0 | 0 | 0 | 0 | 0 |
| *Ziziphus spina-christi* | 0.95 | 0.52 | 0.41 | 1.89 | 1.48 | 1.52 | 0.35 | 3.36 | 0 | 0 | 0 | 0 |

(PA= protected area, GL=grazing land and FL=farm land) RF=Relative frequency, RD-Relative density; RDO=Relative dominancy and IVI= Important Value Index

Woody species like, *Acacia Senegal* (19.01), *Acacia tortilis* (17.75) and *Acacia brevispica* (17.75) with the highest IVI at the GL. This also indicate that these species are the most ecologically significant species in GL. Woody species like, *Barbeya oleoides* (0.75), *Withania somnifera* (0.78) and *Jasminum abyssinicum* (0.80) had a lower IVI at the CGL, attracting conservation effort based on the species’ social and ecological values (Table 3).

In the FL site, *Grewia schweinfurthii* (24.51), *Acacia senegal* (22.47) and *Acacia negrii* (21.49) are the top three ecologically important species with having highest IVI values. *Acacia senegal* had similarly the highest IVI in all three study sites. This shows that there is a little bit of similarity of the sites in most of the factors as they are adjacent to each other. The least ecologically significant species based on their IVI value in this site were, *Dracaena ombet* (0.89), *Sansevieria ehrenbergii* (1.49) and *Grewia villosa* (1.56). The least significant species found in all of the three sites are perfectly different. This also indicate that there is high variation of status of anthropogenic disturbance likes, farming and grading among the three disturbance regimes.

### 3.7. Woody Species Distribution by Diameter Class

The DBH class of species in PA and GL sites were classified in to six class. Whereas, in FL continues only up to five diameter class. In PA and GL vegetation, the total DBH class distribution of woody species indicated an inverted J-shape distribution. This shows where the species DBH class distribution was most common in the smaller diameter and vice versa. In this study, 47.23% and 54.14% of total DBH frequency lies between the first two diameter classes in PA and GL respectively. This indicated that there was drawing out of matured and high diameter class trees for by local people for various purposes (fences, farm implements, house construction, and wood fuel) Similarly, Getaneh (2007) and Tefera et al. (2005) reported that local people gathered woody species with DBH>30 cm for construction and charcoal production. However, most DBH frequency (54.29 percent) was restricted between the second and third diameter classes in FL diameter patterns of woody species, indicating that there were a higher number of individuals in the middle diameter classes, but a reduction towards the lower and higher diameter classes (Fig 3).This indicates a poor reproduction capacity of the species in the community.

**Figure 3.**
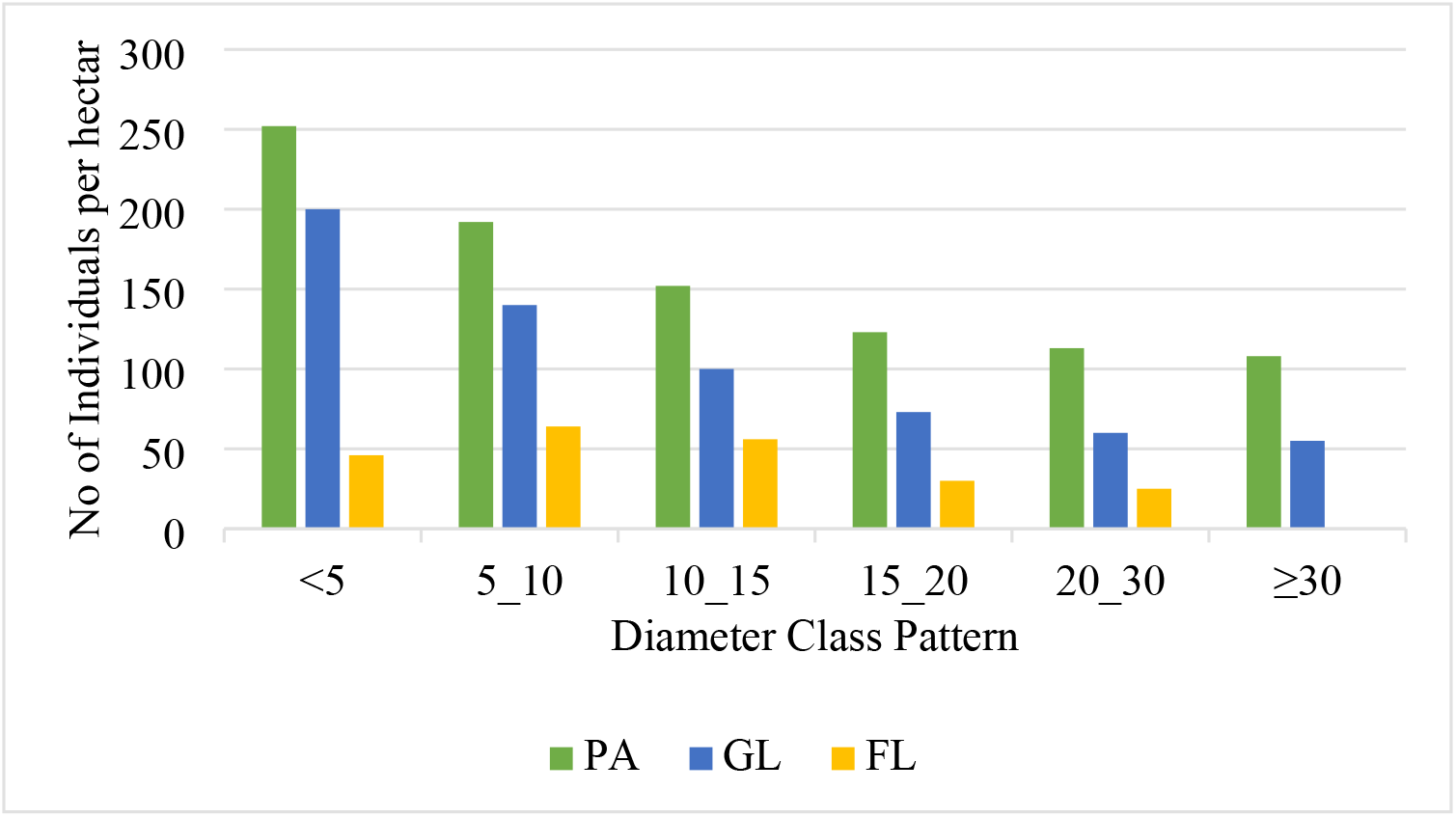
Diameter distribution of species in three land use system (PA= protected area, GL=grazing land and FL=farm land)

**Figure 4.**
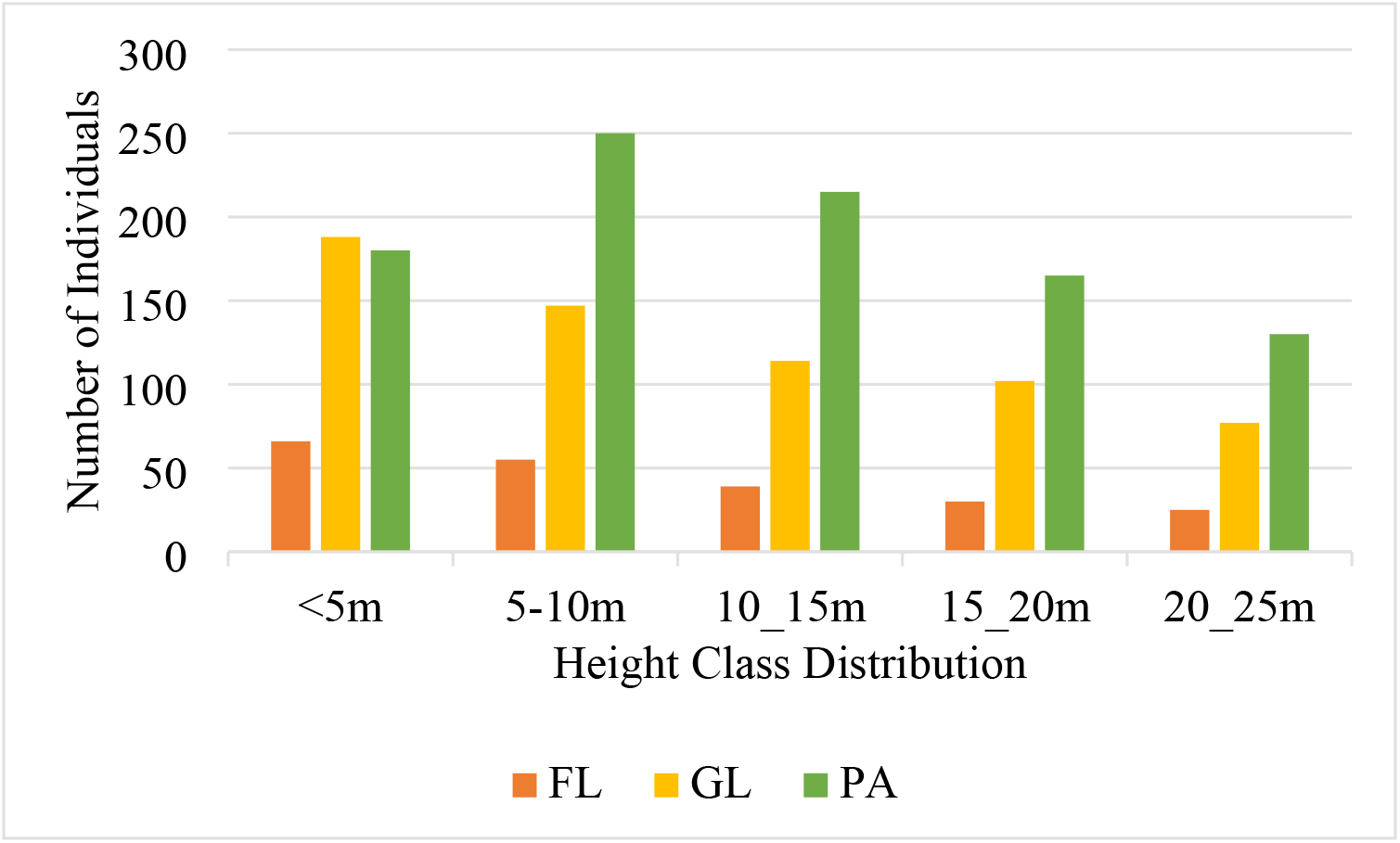
Height class distribution of woody species at the three sites (PA= protected area, GL=grazing land and FL=farm land)

### 3.8. Height Distribution

Height class were classified in to five class in Gola natural vegetation. Unlike that of DBH class, height class distribution shows reversed J-shape pattern in GL and FL sites with gradual decrement towards the largest highest class. This result showed a progress and good regeneration status of population of woody species stability in both GL and FL land use system. However, it reveals irregular pattern in PA, which is dominated by tallest trees and shrubs. most species were attained the medium canopy layer class (5-10 and 10-15m) in PA, this reveals that the number of individuals in the middle classes was the highest, and that the number of individuals in the lower and higher height classes was the lowest. The increased quantity of large-sized individuals in the upper height class in natural vegetation suggests the presence of a good number of grown vegetation species for reproduction (Yeshitela and Bekele, 2003). This argument grips factual for PA of Gola natural vegetation. This is partially because of the absence of large-scale woody exploitation by local dweller.

## 4. Conclusion

The natural vegetation of Gola contains a considerable number of woody species with a moderate diversity. The shrubs species of the Fabaceae family dominated the woody species of Gola’s natural vegetation. However, the majority of species face human pressures in the form of woody cuttings; agricultural development and overgrazing, both of which are regarded ecological and environmental issues, have contributed to the degradation of the vegetation. An investigation of woody species found in PA, GL, and the surrounding FL of Gola natural vegetation revealed the effects of anthropogenic disturbances such as grazing and farming on natural vegetation. The diversity and organization of woody species varied significantly among the three land use categories. In the PA, diverse woody species richness, evenness frequencies, and density were growing, whereas in the adjacent GL to FL, they were declining with a similar set. The population height structure analysis revealed more irregularities in PA, which is dominated by tallest trees and shrubs, with the majority of species reaching the medium canopy layer class, indicating the urgent need for a conservation plan to ensure the long-term viability of woody vegetation resources. To summarize, this kind of thorough vegetation analyses and a comparison analyses aid in determining the influence of conservation techniques on natural vegetation in countries such as Ethiopia, where large-scale conservation initiatives in degraded landscapes are intensively practiced.

## Notes

### Competing Interest Statement

The authors have declared no competing interest.

